# Presenilin-deficient neurons and astrocytes display normal mitochondrial phenotypes

**DOI:** 10.1101/2020.07.15.204255

**Authors:** Sabrina Contino, Céline Vrancx, Nuria Suelves, Devkee M. Vadukul, Valery L. Payen, Serena Stanga, Luc Bertrand, Pascal Kienlen-Campard

**Affiliations:** CEMO-Alzheimer Research Group, Institute of Neuroscience, Université catholique de Louvain, Brussels, Belgium; Laboratory of Advanced Drug Delivery and Biomaterial (ADDB), Louvain Drug Research Institute (LDRI), Université catholique de Louvain, Brussels, Belgium; Neuroscience Institute Cavalieri Ottolenghi, Department of Neuroscience, University of Torino, Torino 10126, Italy; Pole of Cardiovascular Research, Institute of Experimental and clinical Research, Université catholique de Louvain, Brussels, Belgium

**Keywords:** Presenilins, Alzheimer’s disease, OXPHOS, Mitochondria, Astrocyte, Neuron

## Abstract

Presenilins 1 and 2 (PS1 and PS2) are predominantly known as the catalytic subunits of the γ-secretase complex which generates the amyloid-β (Aβ) peptide, the major constituent of the senile plaques found in the brain of Alzheimer’s disease (AD) patients. Apart from their role in γ-secretase activity, a growing number of cellular functions have been recently attributed to PSs. They are involved in synaptic transmission, endo-lysosomal function and calcium homeostasis. PSs were also found to be enriched in mitochondria-associated membranes (MAMs) where mitochondria and endoplasmic reticulum (ER) interact. PS2 was more specifically reported to regulate calcium shuttling between the ER and mitochondria by controlling the formation of functional MAMs through its interaction with the Mitofusin2 protein. We have previously demonstrated that the absence of PS2 (PS2KO) alters mitochondrial morphology and function. Indeed, a PS2KO cell line showed reduced mitochondrial respiration along with disrupted mitochondrial cristae and increased glycolysis. This phenotype is restored by the stable re-expression of human PS2. Still, all these results were obtained in immortalized Mouse Embryonic Fibroblasts (MEF) and one bottom-line question is to know whether these observations hold true for the Central Nervous System (CNS) cells, and in particular neurons and astrocytes. To that end, we carried out primary PS1KO, PS2KO and PS1/PS2KO (PSdKO) neuronal and astrocyte cultures. All the conditions were obtained in the same litter by crossing PS2 heterozygous and PS1 floxed (PS2^+/−^; PS1^flox/flox^) animals. Indeed, contrary to PS2KO mice, PS1KO are not viable and therefore require the use of the Cre-LoxP system to achieve gene deletion *in vitro*. Strikingly, we did not observe any mitochondrial phenotype in PS1KO, PS2KO or PSdKO primary cultures. Mitochondrial respiration and membrane potential were similar in all models, as were the glycolytic flux and NAD^+^/NADH ratio. We further investigated the discrepancies between these results and the ones previously reported in the MEF PS2KO cell line by analyzing PS2KO primary fibroblasts. No mitochondrial dysfunction was observed in this model, in line with observations in PS2KO primary neurons and astrocytes. These results indicate that the mitochondrial phenotype observed in immortalized PS2-deficient cell lines cannot be extrapolated to primary neurons, astrocytes and even to primary fibroblasts. The PS-dependent mitochondrial phenotype reported so far might be the consequence of a cell immortalization process and, therefore, should be critically reconsidered regarding its relevance to AD.

## 1. Introduction

Alzheimer’s disease (AD) is the most prevailing age-related neurodegenerative disease. Its cost and impending rise owed to societal aging makes it a serious social concern and a critical public health burden.

ThIS pathology is characterized by the appearance and spreading of two typical lesions in the brain: senile plaques that are extracellular deposits of the amyloid-β peptide (Aβ), and neurofibrillary tangles (NFTs) that are intracellular aggregates of hyperphosphorylated tau protein (Serrano-Pozo, Frosch, Masliah, & Hyman, 2011). The most admitted comprehensive hypothesis for the onset and development of AD is the amyloid cascade hypothesis (ACH) (Hardy, 2006). It postulates that changes in Aβ production, accumulation or clearance are the triggering events that induce the formation of NFTs, all of which are considered to be responsible for neurodegeneration and dementia. Many efforts have been undertaken to better understand the physio-pathological processes responsible for the altered production of Aβ. They led to the identification of the Amyloid Precursor Protein (APP), the precursor of the Aβ peptide, and the Presenilin proteins (PS1 and PS2) which are the catalytic subunits of the γ-secretase that releases Aβ upon amyloidogenic processing (Goate et al., 1991; Rogaev et al., 1995; Sherrington et al., 1995). Still, clinical trials targeting the amyloid deposition have failed so far to improve cognitive deficits, even though amyloid load can be reduced under certain conditions (Ceyzeriat et al., 2020). Amyloid deposition appears to take place very early in pre- clinical stages and might correlate to the brain dysfunction that appears prior to clinical symptoms. Metabolic aspects of the pathology are widely studied in that context. In imaging studies, the ^18^F-FDG-PET scan is an established imaging biomarker for neuronal dysfunction in the diagnostic workup of AD-patients (Herholz, 2012; Mosconi et al., 2010; Shivamurthy, Tahari, Marcus, & Subramaniam, 2015). This technique underlines that hypometabolism and brain atrophy appear before clinical symptoms in patients. Mitochondria play a central role in cellular metabolism, and mitochondrial function is known to be particularly affected in the disease (Garcia-Escudero, Martin-Maestro, Perry, & Avila, 2013). Studies have put forth mitochondrial dysfunctions as an early cause of AD, suggesting a “mitochondrial cascade hypothesis” (Stanga, Caretto, Boido, & Vercelli, 2020), while others propose that these are a consequence of other pathological processes such as amyloid deposition (Pagani & Eckert, 2011; Swerdlow, 2018; Swerdlow, Burns, & Khan, 2014). Despite this controversy, there is little doubt that mitochondrial dysfunction contributes to AD pathogenesis, as evidenced in animal models of AD (Dixit, Fessel, & Harrison, 2017; Long & Holtzman, 2019; Yao et al., 2009) and samples from AD patients (Adav, Park, & Sze, 2019; Martin-Maestro et al., 2017; Wang et al., 2009). The major consequences of mitochondrial dysfunction are: increased Reactive oxygen species (ROS) production (Dixit et al., 2017), impaired balance of fusion/fission with altered morphology (Wang et al., 2009), decreased oxidative capacity and decreased motility (Correia, Perry, & Moreira, 2016).

These mitochondrial dysfunctions were associated to functional changes of the major AD proteins, namely APP, Tau and PSs (Garcia-Escudero et al., 2013). However, only Presenilin 1 and 2 (PS1 and PS2) were clearly shown to be involved in mitochondrial function. PS1 and PS2 are encoded by two homologous genes, *PSEN1* and *PSEN2*, respectively. Mutations in *PSEN* genes are responsible for inherited forms of AD (FAD) (Hardy, 2006). Apart from their involvement in Aβ production, functions attributed to PSs can be divided into two categories: γ-secretase-dependent or γ-secretase-independent (Vetrivel, Zhang, Xu, & Thinakaran, 2006; Zhang, Zhang, Cai, & Song, 2013). Regarding the γ-secretase independent function of PSs, they are known to be involved in synaptic transmission, endosome-lysosome trafficking, Wnt signaling and, calcium homeostasis which has drawn the greatest research interest. The involvement of PSs in calcium homeostasis has been widely demonstrated. PSs could interact with Inositol trisphosphate receptor (IP3R) (Cheung et al., 2008), sarco-/endoplasmic reticulum (ER) Ca2^+^ ATPase (SERCA) or Ryanodine receptors (RyR) (Green et al., 2008; Wu, Yamaguchi, Lai, & Shen, 2013) to regulate intracellular calcium signaling. PSs were also suggested to directly act as a low-conductance, passive ER Ca2^+^ channel (Nelson et al., 2007). Finally, PSs could regulate the calcium crosstalk between the mitochondria and the ER by regulating their apposition through a particular domain called mitochondria-associated membranes (MAMs) (Filadi et al., 2016). Indeed, studies have shown that PSs are enriched in MAMs, which are lipid-raft-like structures. MAMs are considered as functional compartments (Area-Gomez et al., 2009) due to their implication in cellular pathways such as inflammation, mitophagy and lipid production (Filadi, Theurey, & Pizzo, 2017). PS2 was found to physically interact with Mitofusin 2 to regulate MAMs organization and calcium shuttling in Mouse Embryonic Fibroblasts (MEF). Disruption of this interaction and of the consecutive calcium crosstalk was reported in a PS2KO MEF cell line (Zampese et al., 2011). Importantly, impaired calcium influx from the ER to mitochondria is implicated in the regulation of the oxidative phosphorylation (OXPHOS) and can lead to mitochondrial defects. Using the same MEF cell lines, we observed an altered mitochondrial phenotype related to the absence of PS2 and not PS1 (Contino et al., 2017). PS2 deficiency results in defective mitochondrial cristae correlated to an impaired OXPHOS capacity and a modified redox state (NAD^+^/NADH ratio). This was compensated by an increased glycolytic capacity, sustaining a stable ADP/ATP ratio. Together with other studies (Correia et al., 2016; Dixit et al., 2017; Pagani & Eckert, 2011; Wang et al., 2009), this led to the hypothesis that PS-dependent mitochondrial dysfunction could represent a major pathway in AD pathogenesis.

In this context, we investigated for the first-time mitochondrial activity in primary cultures of neurons and astrocytes, which are more relevant cellular models to AD. We performed primary cultures of cells expressing both PSs (wild-type, WT), one of them (PS single-knockout, referred to as PS1KO and PS2KO) or none (PSs double-knockout, referred to as PSdKO). Our aim was to investigate (i) if mitochondrial-related deficits appear in the absence of PSs in these cells (ii) and, in so, to identify which PS is involved in this phenotype and the possible mechanisms underpinning neuronal/astrocytic dysfunction. Surprisingly, we did not find any metabolic deficit in the different PS knockout primary nerve cells, even when both PSs were absent (PSdKO). This suggests that the PS2-related mitochondrial deficiency reported in the MEF cell line does not occur in neurons or astrocytes. We further studied the mitochondrial activity in PS2KO primary MEF, and did not observe any mitochondrial defect. With all these evidences, we postulate that PS-dependent mitochondrial alterations appear only in immortalized cells, and may provide information about PS-dependent cancer processes much more than about mitochondrial dysfunction in AD.

## 2. Material and methods

### 2.1. Animal models

Presenilin 2 knockout (KO) (#005617) (Herreman et al., 1999) and Presenilin 1 floxed (#004825) (Yu, Kessler, & Shen, 2000) mice, both in a C57BL6 background, were obtained from Jackson Laboratories (Bar harbor, USA). All animal procedures and experiments were approved and performed in agreement with the UCLouvain animal care committee’s regulations (code number 2016/UCL/MD/016). Animals were housed on a 12 h light/dark cycle in standard animal care facility with access to water and food *ad libitum*.

For primary cell cultures, generation of the different genotypes in the same litter were obtained from crossing PS2 heterozygous and PS1 floxed (PS2^+/−^; PS1^flox/flox^) animals followed or not by viral transduction. Indeed, *PSEN1 gene* deletion was achieved by viral transduction of Cre recombinase in floxed PS1 primary cultures. Lentivirus expressing GFP (mock control) or CRE-GFP were used for transduction at DIV 1 (one day of *in vitro* culture) for neurons and DIV 17 for astrocytes. Following genotyping and infection we could obtain in the same litter: control non-infected **Ct** (PS2^+/+^; PS1^flox/flox^, non-infected), **PS2KO** (PS2^−/−^; PS1^flox/flox^, non-infected); control infected named **Mock** (PS2^+/+^; PS1^flox/flox^, infected with GFP); **PS1KO** (PS2^+/+^; PS1^flox/flox^, infected with CRE-GFP) and **PSdKO** (PS2^−/−^; PS1^flox/flox^, infected with CRE-GFP).

### 2.2. Primary neuronal cultures

Primary cultures of neurons were performed as previously described (Opsomer et al., 2020) on E17 mouse embryos. By dissection, cortices and hippocampi were isolated on ice cold HBSS (Thermo Fisher Scientific, Waltham, USA) and meninges were removed. Tissues were then dissociated by pipetting up and down 15 times with a glass pipette in HBSS glucose 5mM medium. Dissociation was repeated 10 times with a flame narrowed tip glass pipette. After sedimentation for 5 min, the supernatant containing the neurons was settled on a bed of 4 ml of Fetal Bovin Serum (FBS) and centrifuged at 1,000 × *g* for 10 min. The pellet was resuspended in Neurobasal^®^ medium enriched with 1 mM L-glutamine, 5 mM glucose and 2% v/v B-27^®^ supplement medium. Cells were plated at 200,000 cells/cm^2^ on pre-coated poly-L-lysine dishes and cultured (37°C, 5% CO_2_ and humidified atmosphere). Half media changes were performed every 2 days and neurons were cultured for 11 days (DIV 11) before being utilized for experiments. To get PS1KO and PSdKO, neurons were infected at DIV 1 and all media were changed at DIV 2.

### 2.3. Primary astrocyte cultures

For primary astrocyte cultures, the brains of mouse pups aged 2 days were dissected to isolate cortices on ice cold HBSS. Tissues were triturated 15 times with glass pipette and 10 times with flame narrowed tip glass. Tubes were centrifuged at 1,000 × *g* for 5 min. Pellets were resuspended in HBSS and centrifuged at 1,700 × *g* for 20 min on a 30% Percoll gradient. Astrocytes were collected from the interphase, washed in 10 ml of HBSS and centrifuged for 5 min at 1,500 × *g*. Pellets were resuspended and plated in DMEM-glutaMAX medium (Thermo Fisher Scientific, Waltham, USA) supplemented with 10% FBS (Biowest, Nuaillé, France), 50 mg/ml penicillin–streptomycin and 50 mg/ml fungizone. Cells were left to proliferate in flasks for 15 days at 37°C and 5% CO_2_, and media were changed every 4-5 days. After 15 days, astrocytes were plated and further cultured in DMEM-glutaMAX with 10% FBS. Two days later, transduction with lentivirus (if necessary) and differentiation was induced by reducing the concentration of FBS to 3% for 7 days before performing experiments (DIV7).

### 2.4. Primary and immortalized mouse embryonic fibroblasts (MEF)

Immortalized MEF were cultured as previously described (Contino et al., 2017). Primary cultures of MEF were performed on E16 embryos. Chest skin was isolated on ice and then grinded into pieces with a blade. These pieces were dissociated and incubated twice at 37°C for 10 min in trypsin (Life Technologies, Carlsbad, USA). DMEM low glucose (5.5 mM) (Sigma-Aldrich, St Louis, USA) enriched with 10% FBS and 1% pen/strep, was added for an incubation of 5 min at room temperature (RT). Supernatants were collected and centrifuged at 1000 × *g* for 5 min at RT. Pellets were resuspended in 10 ml of DMEM medium and plated in petri dishes. Once at confluence, cells were plated for experiments.

### 2.5. Lentiviral particles

Lentiviral particles expressing CRE recombinase were used to delete floxed *PSEN1*. Plasmids pCMV-GFP (Mock) and pCMV-CRE-GFP for lentiviral production were purchased from Cellomics Technology (Halethorpe, USA). Amplification and purification of the different plasmids were performed using the Plasmid Midi kit (Qiagen, Hilden, Germany). Production was carried out by transfecting HEK293-T cells in 10 cm dishes (2×10^6^ cells/dish) with both CRE-GFP or GFP vectors, pMD2.G (Addgene#12259) and pCMV-dR8.2 (Addgene#12263). 48 h after transfection, cells were harvested by flushing with the medium and centrifuged at 1,500 × *g* for 10 min at 4°C. The supernatant was filtered with Acrodisc^®^ 0.45 μm filters (Pall, NYC, USA). 1/3 (v/v) of LentiX™ Concentrator reagent (Clontech, Mountain View, USA) was added and incubated overnight (o/n). After centrifugation at 1,500 × *g* for 45 min at 4°C, the pellet was resuspended in 20 μl per dish of DMEM without serum and stored at −80°C. 10 μl of concentrated virus was used to infect 1,600,000 neurons or 300,000 astrocytes.

### 2.6. Western blotting (WB)

WB was performed on cell lysates obtained by harvesting cells with lysis buffer (Tris 125 mM pH 6.8, 4% sodium dodecyl sulfate, 20% glycerol) with Complete Protease Inhibitor Cocktail (Roche, Basel, Switzerland) and sonicating, as previously described (Stanga et al., 2016). Protein concentration was determined using the BCA Protein assay kit (Pierce, Rockford, USA). 15 μg of total proteins diluted in lysis buffer with NuPage Sample Buffer and 0.5M DTT was heated at 70 °C (except for the detection of OXPHOS subunits, heated at 37°C). Samples were loaded and separated onto NuPAGE^®^ 4-12% Bis-Tris gel (Life Technologies, Carlsbad, USA) with a NuPAGE^®^ MES SDS running buffer (Life Technologies, Carlsbad, USA). Proteins were then transferred onto an Amersham™ nitrocellulose membrane (Little Chalfont, UK) with NuPAGE^®^ transfer buffer (Life Technologies, Carlsbad, USA) for 2 h at 30 V. After blocking with nonfat dry milk (Darmstadt, Germany) (5% in PBS and 0.05% Tween-20) for a minimum of 30 min, the membrane was incubated o/n with the primary antibody diluted in PBS 0.05% Tween. Secondary antibody coupled to horse radish peroxidase was incubated 1 h at RT. Detection with ECL PerkinElmer^®^ (Waltham, USA) was performed.

Antibodies used and their dilutions are the following: Anti-TOM20 (1:1,000; Proteintech, Rosemont, USA); Anti-OXPHOS Cocktail (1:1,000; Abcam, Cambridge, United Kingdom); Anti-Presenilin 1 (1:1,000; Cell Signaling, Danvers, USA);); Anti-Presenilin 2 (1:1,000; Cell Signaling, Danvers, USA); Anti-Tubulin (1:3000, Abcam, Cambridge, United Kingdom); Anti-Actin (1:3,000; Abcam, Cambridge, United Kingdom); Anti-mouse (1:10,000; GE Healthcare, Little Chalfont, United Kingdom); Anti-rabbit (1:10,000; GE Healthcare, Little Chalfont, United Kingdom).

### 2.7. Immunocytochemistry (ICC)

ICC were performed as previously described (Stanga et al., 2018). Cells were seeded on pre-coated poly-L-lysine coverslips in 24 well plate at the density of 200,000/well neurons or 50,000/well for astrocytes. Cells were fixed with PBS/paraformaldehyde 4% for 10 min and rinsed three times for 5 min with PBS. Cells were then permeabilized with a solution of PBS/ Triton 0.3% for 30 min and non-specific sites were blocked with PBS/Triton 0.3%/ FBS 5% for 30 min. Primary antibodies diluted in the blocking solution were incubated o/n at 4°C. After 3 washes of 10 min with PBS, cells were incubated with DAPI and secondary antibodies diluted in the blocking solution. Dilutions of the antibodies were as follows: chicken anti glial-fibrillary acidic protein (GFAP2; 1:1,1000; Abcam, Cambridge, United Kingdom); mouse anti-microtubule associated protein (MAP2; 1:1,000; Sigma‐Aldrich; St Louis, USA); DAPI (1:2,000; Sigma‐Aldrich; St Louis, USA); Alexa 488 anti-rabbit (1:500; Life Technologies, Carlsbad, USA). Images were acquired on EVOS FL Auto microscope (Invitrogen) with RFP (Alexa Fluor 554), and CY5 (Alexa Fluor 647) EVOS LED light cubes and analyzed with ImageJ software.

### 2.8. Mitochondrial membrane potential (ΔΨ)

Fluorescent cationic probe tetramethylrhodamine methyl ester (TMRM) (Sigma-Aldrich, St Louis, USA) was used to evaluate the ΔΨ. As a control, we used the uncoupling agent Carbonyl cyanide-4-(trifluoromethoxy) phenylhydrazone (FCCP) (Sigma-Aldrich, St Louis, USA). Cells were plated in 96 well plates at a density of 60,000/well for neurons or 15,000/well for astrocytes. Cells were incubated for 30 min at 37°C with TMRM (30 nM) with or without FCCP (10 μM) diluted in KREBS medium. Cells were then washed with KREBS solution and fluorescence was read with the plate reader VICTOR^®^ Multilabel Plate Reader (PerkinElmer). Data were normalized to the total amount of protein measured by the Bradford assay kit (Bio-Rad Laboratories, California, USA).

### 2.9. Mitochondrial oxygen consumption

Oxygen consumption rate (OCR) was measured with the Seahorse XF96 bioenergetic analyzer (Seahorse Bioscience; Massachusetts, USA). Cells were seeded in a Seahorse 96 well plate at different several densities (60,000/well for neurons, 15,000/well for astrocytes or 20,000/well for MEF). To analyze of the effect of the inhibition of γ-secretase activity on OCR, cells were treated with N‐[N‐(3,5‐difluorophenacetyl)‐L‐alanyl]‐sphenylglycine t‐butyl ester (DAPT) (Calbiochem, Camarillo, CA, USA) 24 h before performing the experiment. Once differentiation was completed, the medium was exchanged with the conditional medium (culture medium without sodium bicarbonate and FBS) and incubated without CO_2_ at 37°C for 1 h. Inhibitors targeting the different mitochondrial complexes (Cell Mito Stress Test kit, Seahorse Bioscience) were added automatically and sequentially to the cells during the experiment to measure the basal respiration, the coupling and the spare respiratory capacity. The sequence of the inhibitors used was: Oligomycin (1 μM); FCCP (0.5 μM for neurons and 1 μM for astrocytes and MEF); Rotenone and antimycin A (0.5 μM). Results were normalized to the total amount of protein measured by the Bradford assay kit (Bio-Rad Laboratories, California, USA).

### 2.10. NAD^+^/NADH ratio

NAD^+^/NADH ratio was measured with the bioluminescent NAD+/NADH-Glo assay kit (Promega, Wisconsin, USA) according the manufacturer’s instructions. Cells were seeded in a 96 well plate at different densities (60,000/well for neurons, 10,000/well for astrocytes or 20,000/well for MEF). Once the differentiation was completed, cells were rinsed with PBS and then lysed with the basis solution 1 (% dodecyltrimethylammonium bromide (DTAB). Samples were split for a basic or acid treatment and were heated at 60°C for 15 min. The reduced form was decomposed in the acidic solution and the oxidized form was selectively decomposed in the basic solution. After neutralization, samples were mixed with NAD^+^/NADH-Glo™ detection reagent and incubated for 45 min to induce reaction. Luminescence was read on the GloMax^®^ 96-well plate luminometer (Promega, Wisconsin, USA).

### 2.11. Glycolytic flux

Glycolytic rate was analyzed by the measurement of the detritiation rate of [3-^3^H] glucose as previously described (Contino et al.,2017). Briefly, cells were seeded in a 12 well plate at density of 800,000/well for neurons, 100,000/well for astrocytes or 200,000/well for MEF. Tritiated glucose (0.2 μCi/ml; Perkin-Elmer; Massachussets, USA) was added to the medium (including 5.5 mM glucose) for 30 min. After medium collection, the tritiated water resulting from detritiated glucose was separated from the non-transported tritiated glucose by column chromatography and measured with the Tri Carb 2,810 liquid scintillation analyzer (Perkin Elmer; Massachussets, USA) as described previously (Marsin, Bouzin, Bertrand, & Hue, 2002). Data were normalized to the total amount of protein measured by BCA assay (Thermo Scientific, Rockford, USA).

### 2.12. ATP level

ATP level was measured with the bioluminescent ATPlite^®^ assay kit (Promega, Wisconsin, USA) according the manufacturer’s instructions. Cells were seeded in a 96 well plate at density of 60,000/well for neurons or 10,000/well for astrocytes. On the day of the experiment, medium was replaced by 75 μl of fresh medium for 30 min at room temperature. 75 μl of kit’s solution was then added and after 10 min of incubation, luminescence was read on the GloMax^®^ 96-well plate luminometer (Promega, Wisconsin, USA).

### 2.13. Statistical analysis

Number of samples are indicated in figure legends with “n=” and number of independent experiments with “N=”. GraphPad Prism software (GraphPad Software, La Jolla, CA, USA) was used to analyze the data and perform the statistical analyses. Normality was assessed with Shapiro Wilk test (GraphPad Prism). Parametric test (Student’s *t* test, ANOVA and Tukey’s multiple comparison test) was applied if the data followed normal distribution. Otherwise, non-parametric test (Mann-Whitney test, Kruskal-Wallis and Dunn’s multiple comparison test) was used. Significance is indicated as: \**p* < 0.05, \*\**p* < 0.01, \*\*\**p* < 0.001, ****p < 0.0001.

## 3. Results

### 3.1. Model characterization and efficient generation of PS1KO, PS2KO, PSdKO primary nerve cells

Metabolic analyses were performed on primary neurons and astrocytes taken respectively at embryonic day 17 (E17) and at postnatal day 2 (P2). Neurons were cultured for one day (DIV1) and astrocytes for seventeen days (DIV17) prior to lentiviral transduction and further maintained in culture until DIV11 for neurons and DIV24 for astrocyte. The experimental workflow is shown in Figure 1A. These two major brain cell types are known to have different bioenergetic and metabolic profiles: neurons are more oxidative and astrocytes more glycolytic. The different culture times (DIV) were chosen in order to obtain differentiated and active primary neurons and astrocytes (Charlesworth, Cotterill, Morton, Grant, & Eglen, 2015; Schildge, Bohrer, Beck, & Schachtrup, 2013). Generation of the different genotypes in the same litter were obtained from crossing PS2 heterozygous; PS1 floxed (PS2^+/−^; PS1^flox/flox^) animals followed where necessary by viral transduction. Indeed, PS2 knockout (PS2^−/−^; PS2KO) cells are obtained from the viable PS2 full KO mice (Herreman et al., 1999) but PS1KO and PSdKO mice present a lethal phenotype at E17 and E12 respectively (Donoviel et al., 1999; Shen et al., 1997). Therefore, to obtain PS1KO and PSdKO primary cultures, we achieved *PSEN1* deletion by viral transduction of Cre recombinase in PS1^flox/flox^ primary cultures (Figure 1A). Lentivirus inducing GFP (mock control) or CRE-GFP expression were used for transduction of neurons at DIV1 and OF astrocytes at DIV17. The expression level of PS1 and PS2 was monitored by WB in neuronal and astrocytic cultures (Figure 1B). Cre-mediated gene excision of PS1 was very efficient under these conditions (90% decrease when compared to control (Figure 1B).

**Figure 1:**
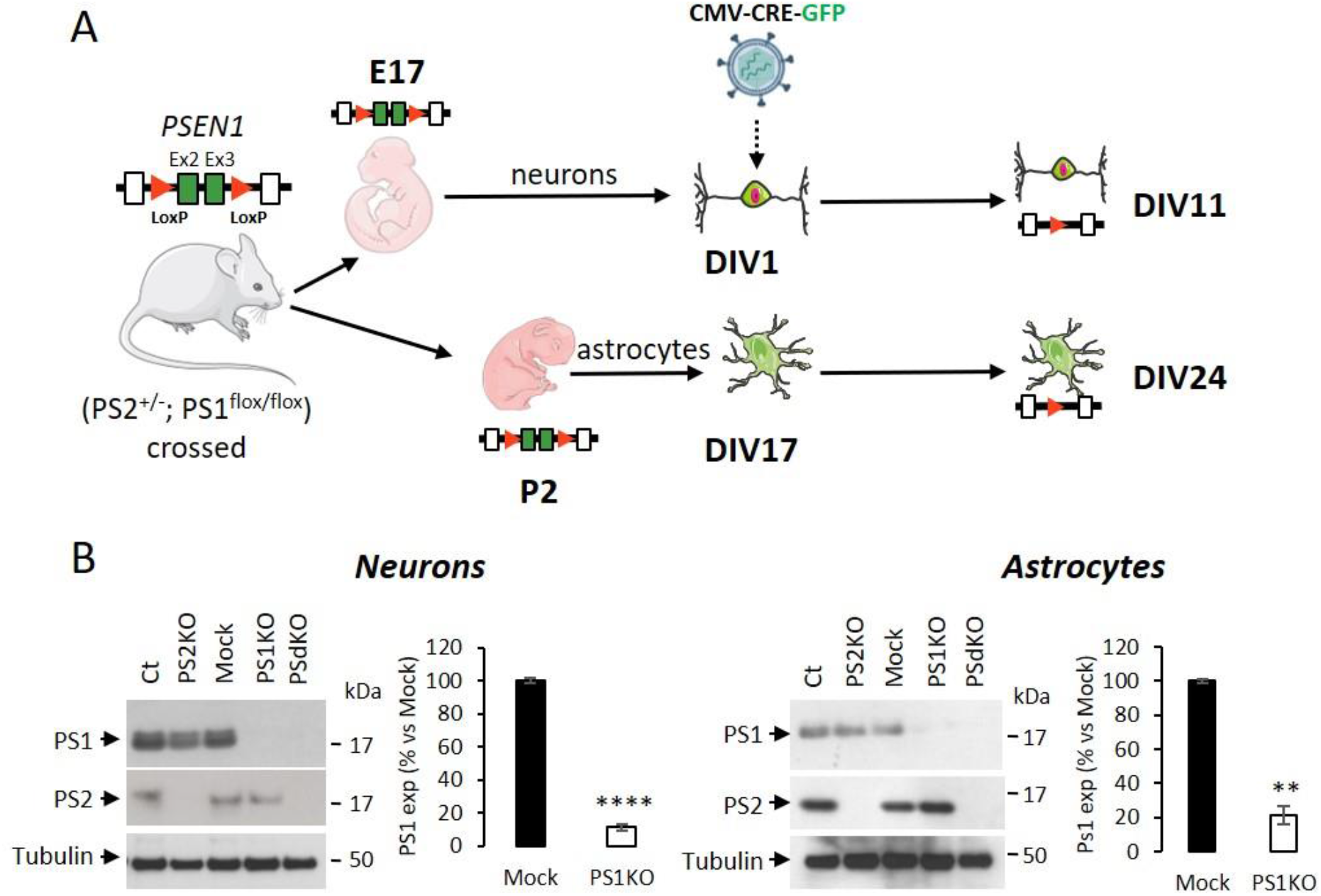
Experimental workflow and model characterization in PS-defective primary nerve cells. **(A)** Workflow description of the experiments with specificity for PSEN1 deletion. Primary neuronal and astrocyte cultures were performed on embryos at day 17 (E17) and postnatal pups at day 2 after birth (P2), respectively. Generation of the different genotypes in the same litter were obtained from crossing PS2 heterozygous and PS1 floxed (PS2^+/−^; PS1^flox/flox^) animals followed where necessary by viral transduction. At DIV1 for neurons and DIV17 for astrocytes, cells were either non-infected (Ct and PS2KO) or infected with GFP (Mock) or CRE-GFP to induce PSEN1 gene deletion (PS1KO and PSdKO). Experiments were performed after 11 days for neurons and 24 days (including 7 days of differentiation) for astrocytes. Ex2 and Ex3 = PSEN1 exons 2 and 3 respectively. **(B)** Representative WBs showing PS1 and PS2 expression profiles in primary neuronal and astrocyte cultures. Actin was used as loading control. Quantification of PS1 expression in total cell lysates is measured by WB. Results (means ± sem) are given as percentage of control cells (Mock); ****p < 0.0001; **p < 0.01; Student’s t test for neurons results and Mann-Whitney test for astrocytes results (N = 6).

In order to characterize our model, we checked the overall morphology of the various PS-deficient cells at DIV11 (neurons) and DIV24 (astrocytes) by immunostaining cells with astrocyte (GFAP) and neuronal (MAP2) markers (Figure 2). Protoplasmic astrocytes cultures were pure while some activated astrocytes were present in neuronal cultures (+/− 10(%). There were no apparent differences in terms of morphology or growth between Ct, mock, PS1KO, PS2KO and PSdKO cells.

**Figure 2:**
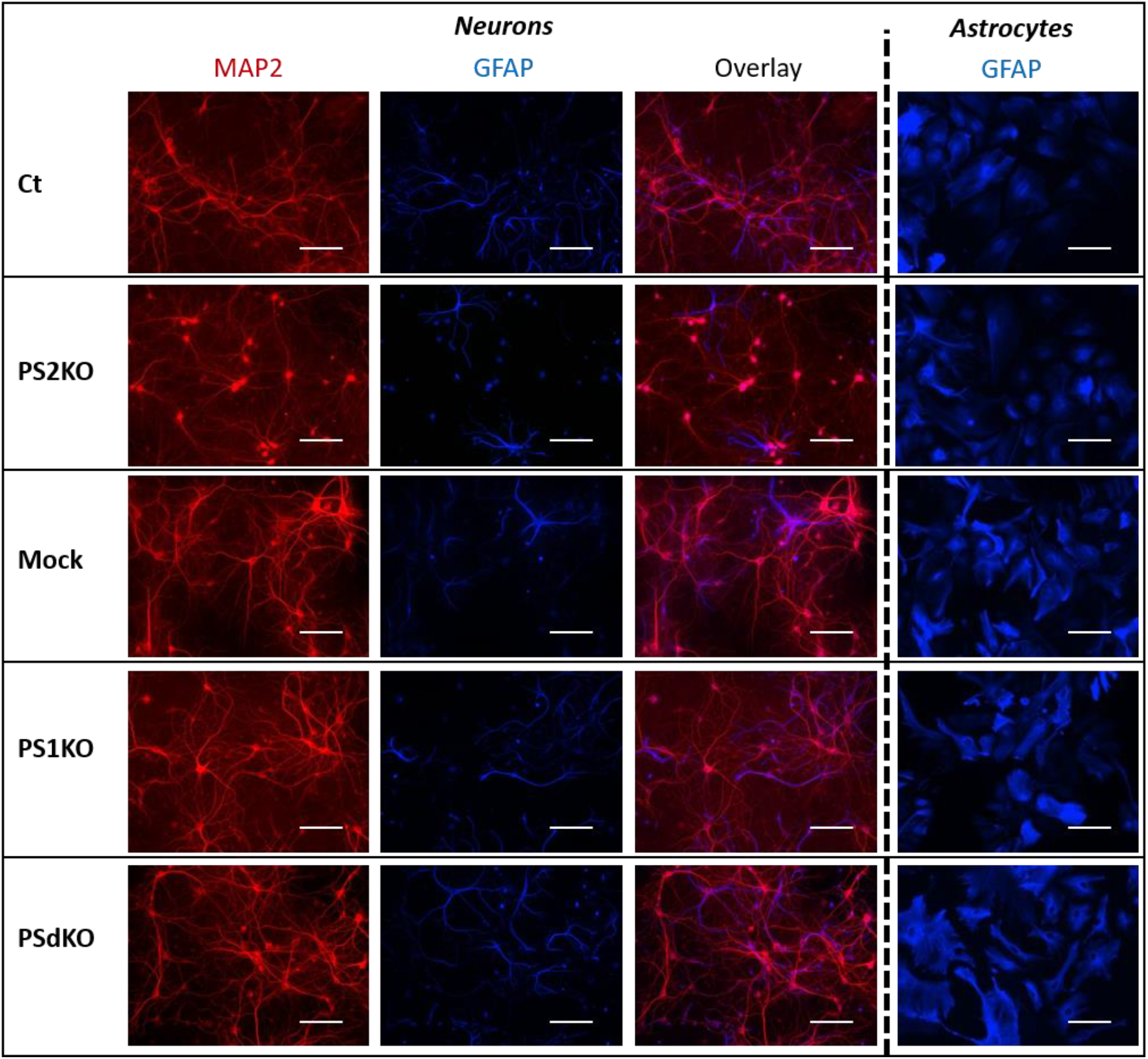
Morphology of PS-defective primary neuronal and astrocyte cultures. On the left side, a primary culture of neurons at DIV11 immuno-stained with glial-specific protein GFAP (blue) and the neuron-specific protein MAP2 (red) antibodies. On the right side, a primary culture of astrocytes at DIV24 immuno-stained with GFAP (blue antibody). Scale bar= 100μM

### 3.2. Membrane potential (ΔΨ) and OXPHOS complexes expression are not affected in PS-deficient primary nerve cells

As a primary indicator of mitochondrial mass (Whitaker-Menezes et al., 2011), the expression level of the mitochondrial import receptor subunit TOM20 (Figure 3A) was first evaluated in cell lysates from primary neurons and astrocytes. No significant difference in TOM20 levels was observed between the different cell types, suggesting that deletion of PSs did not impact the mitochondrial mass in the cells tested. Mitochondrial membrane potential (ΔΨ) is crucial for energy production and it is the driving force generated by the electron transport chain (ETC) for ATP synthesis. ΔΨ is known to stay stable since its decrease is a strong indication signal of cell death (Uechi, Yoshioka, Morikawa, & Ohta, 2006). In this regard, we evaluated the ΔΨ with the TMRM probe (Figure 3B) and FCCP, an uncoupling agent abolishing ΔΨ, used as a control. No significant differences in ΔΨ were observed between the controls (Ct and Mock) and PS2KO, PS1KO and PSdKO cells, neither in primary neurons or astrocytes. Since ATP synthase might work in reverse to keep ΔΨ stable (Uechi et al., 2006), an ETC defect could still occur. Therefore, we checked the expression of the ETC subunits by WB using a cocktail of antibodies targeting representative subunits of the five mitochondrial complexes (Figure 3C). Our results showed that there were no differences in the expression of any subunit in PS1KO, PS2KO or PSdKO neurons or astrocytes. This was rather unexpected since we previously observed a defect in oxidative phosphorylation (OXPHOS) capacity along with expression changes of the ETC subunits in the PS2KO MEF cell line (Contino et al., 2017).

**Figure 3:**
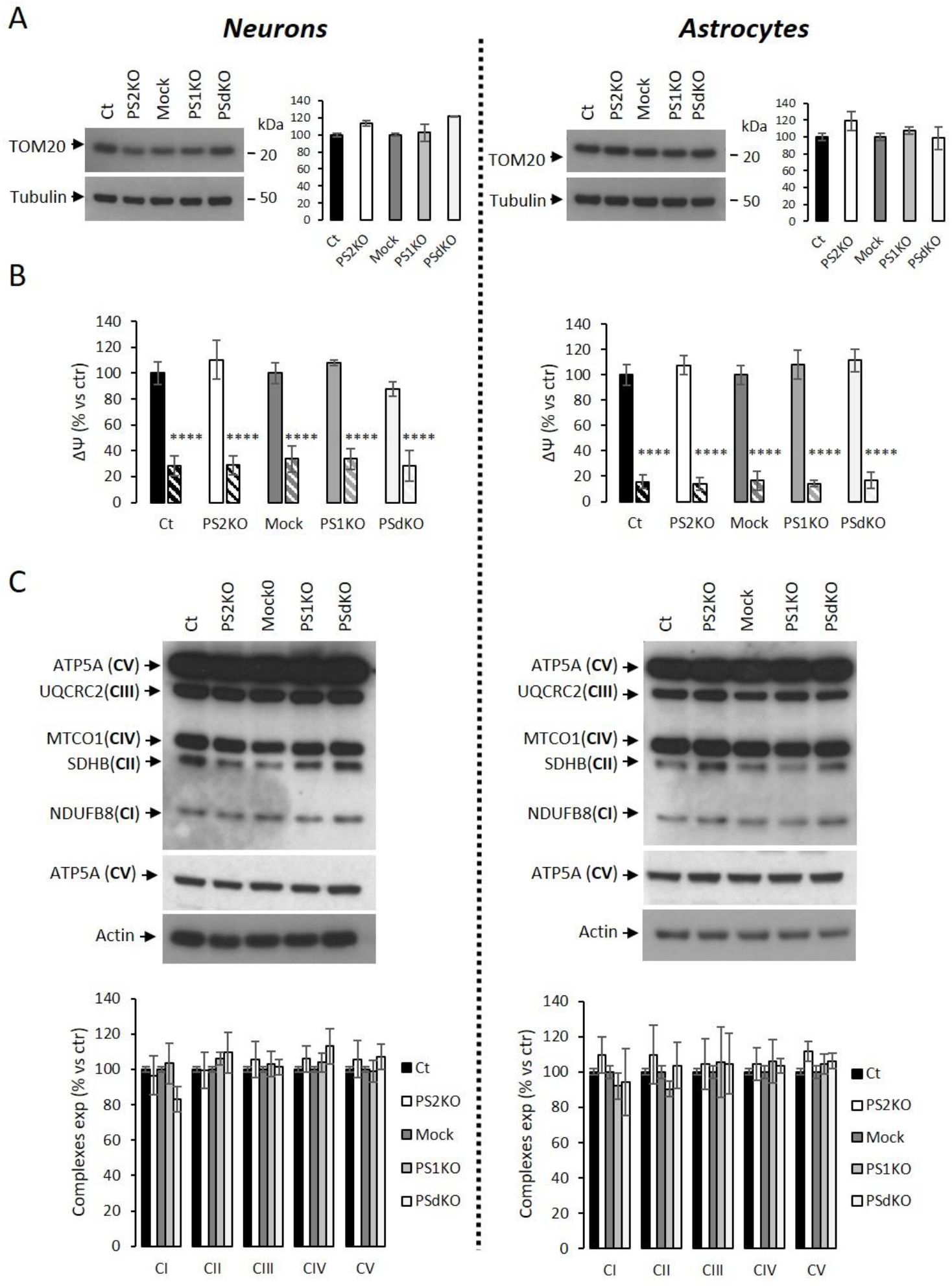
ΔΨ and OXPHOS complexes expression in PS-deficient primary nerve cells. Experiments were performed on primary DIV11 neuronal cultures and primary DIV24 astrocytes cultures. Cell conditions were Ct vs PS2KO, Mock (infection control) vs PS1KO and PSdKO. **(A)** Representative WB and quantification of TOM20 profile expression; in primary neuronal cell lysates (left panel) and in primary astrocyte cell lysates (right panel). Tubulin was used as a loading control. Results (mean ± SEM) are expressed as percentage of the respective control (ctr= Ct or Mock) (min N=2). Kruskal–Wallis test and Dunn’s multiple comparison test. **(B)** ΔΨ was evaluated with the TMRM probe in the presence or absence of FCCP, an uncoupling agent used as control (striped columns). Fluorescence signal was measured with a plate reader and results are expressed as the percentage of the relative mean fluorescence of the respective control cells (ctr =Ct or Mock) (min N=3). ANOVA and Tukey’s multiple comparison test. ****p < 0.0001. **(C)** The expression level of representative protein subunits from each of the five mitochondrial complexes (NDUFB8 for CI; SDHB for CII; UQCRC2 for CIII; MTCO1 for CIV; ATP5A for CV) was evaluated by WB on cell lysates. ATP5A for CV is shown at low (middle panel) and high exposure (upper panel). Actin was used as a loading control. Quantification of the different WBs (means ± sem) are given as percentage of signal measured in the respective control cells (Ct or Mock) (min N= 3). ANOVA and Tukey’s multiple comparison test.

### 3.3. Mitochondrial oxidative phosphorylation and bioenergetics are not affected by the absence of PS1 and/or PS2 in primary neurons and astrocytes

Although if the profile of complexes expression did not show any defect, this biochemical indication does not rule out the hypothesis that OXPHOS could be impaired in PS-deficient primary neurons or astrocytes. Indeed, the cocktail of OXPHOS antibodies used targets only one subunit of each of the massive ETC complex. Hence, we evaluated the activity of the ETC by measuring the overall profile of oxygen consumption rate (OCR) and several related parameters (Figure 4A-C). The parameters measured were basal respiration, coupling (oxygen consumption devoted to ATP synthesis under resting conditions) and spare respiratory capacity (maximal uncoupled rate of respiration minus the basal rate). All these parameters were found to be affected by the absence of PS2 in MEF cell lines as shown in supplemental data Figure 1A. Strikingly, the general OCR measured was similar in the presence (Ct, Mock) or absence of PSs (PS1KO, PS2KO, PSdKO) in primary neurons and astrocytes (Figure 4A-C).

**Figure 4:**
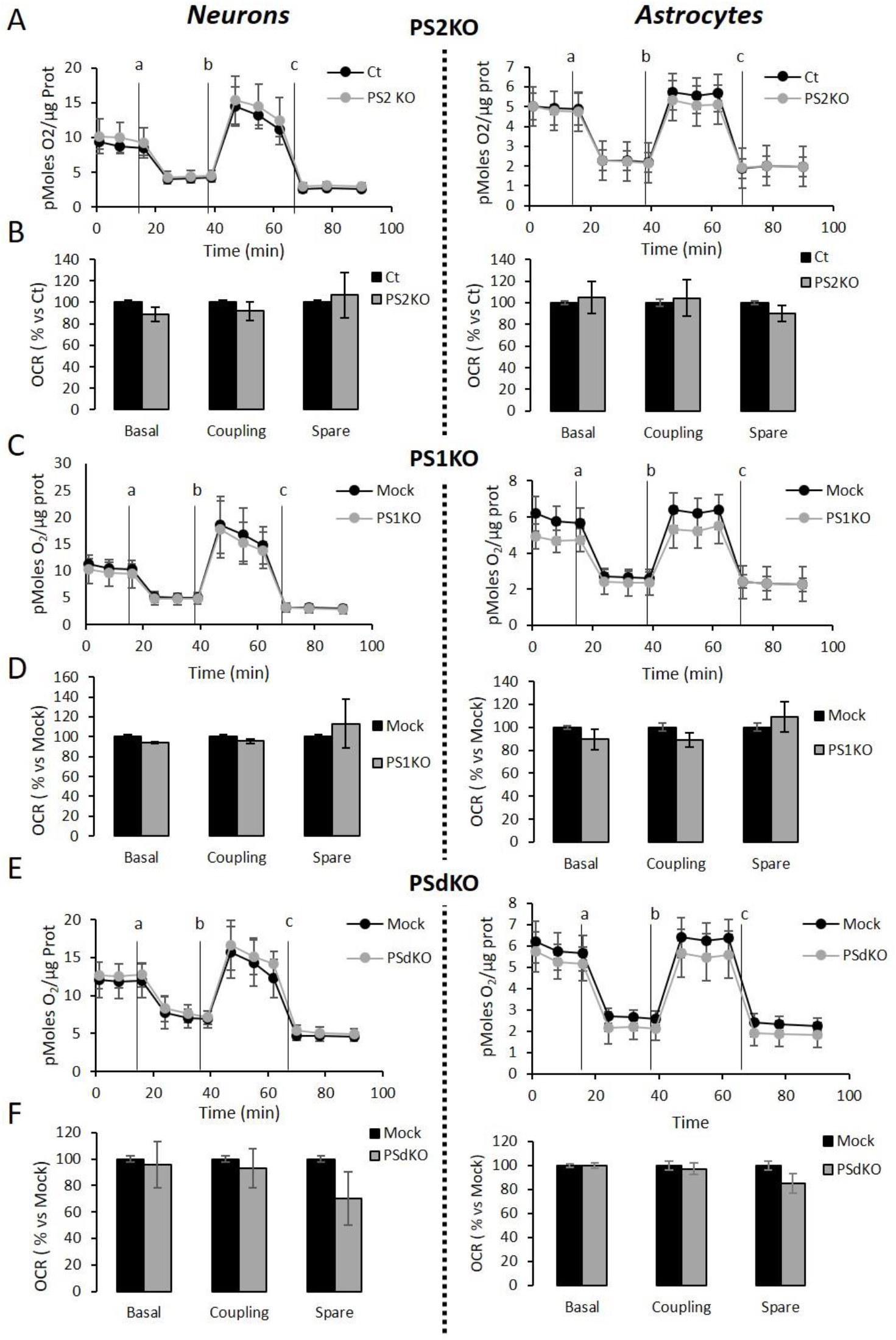
Assessment of the OXPHOS capacity in primary nerve cultures genetically deleted for PSs. Oxygen consumption rate (OCR) was evaluated by using the Seahorse XF96 bioenergetic analyzer. Experiments were carried out in primary DIV11 neuronal cultures (left panel) and in primary DIV24 astrocyte cultures (right panel). Cell conditions were wild type non-infected (Ct) vs PS2KO; control infection (Mock) vs PS1KO and PS1/2 double KO (PSdKO). **(A, C, E)** General profile of the OCR with vertical lines indicating the time point at which the different compounds have been added: a. Oligomycin (CV inhibitor) b. FCCP (ΔΨ uncoupler) c. Rotenone (CI inhibitor) and antimycin A (CIII inhibitor). Values (means ± sem) are given in pmol O_2_/min/μg protein (min N=3). **(B, D, F).** The basal respiration, the coupling ratio and the spare respiratory capacity were calculated according to the Cell Mito Stress Test kit’s recommended protocol. Values (means ± sem) are given as percentage of signal measured in the respective control cells (Ct or Mock) (min N=3). ANOVA and Tukey’s multiple comparison test.

In addition to genetic deletion of PSs, another way to address the unexpected results shown in Figure 4 is to evaluate if accurate inhibition of γ-secretase activity would be impaired OXPHOS. Indeed, PSs are the catalytic core of the γ-secretase complex cleaving more than 90 substrates and their multiple contribution to cellular physiology were shown to be dependent or independent of this function (Zhang et al., 2013). We measured OCR on primary neurons and astrocytes (Figure 5A-B) treated for 24 h with 10 μM of DAPT, a concentration that efficiently blocks γ-secretase activity (Stanga et al., 2018). No changes in OCR were observed in DAPT-treated cells compared to non-treated cells. This confirms that γ-secretase activity is not involved in OXPHOS, which is in line with the facy that the absence of PSs does not affect mitochondrial activity in primary nerve cells.

**Figure 5:**
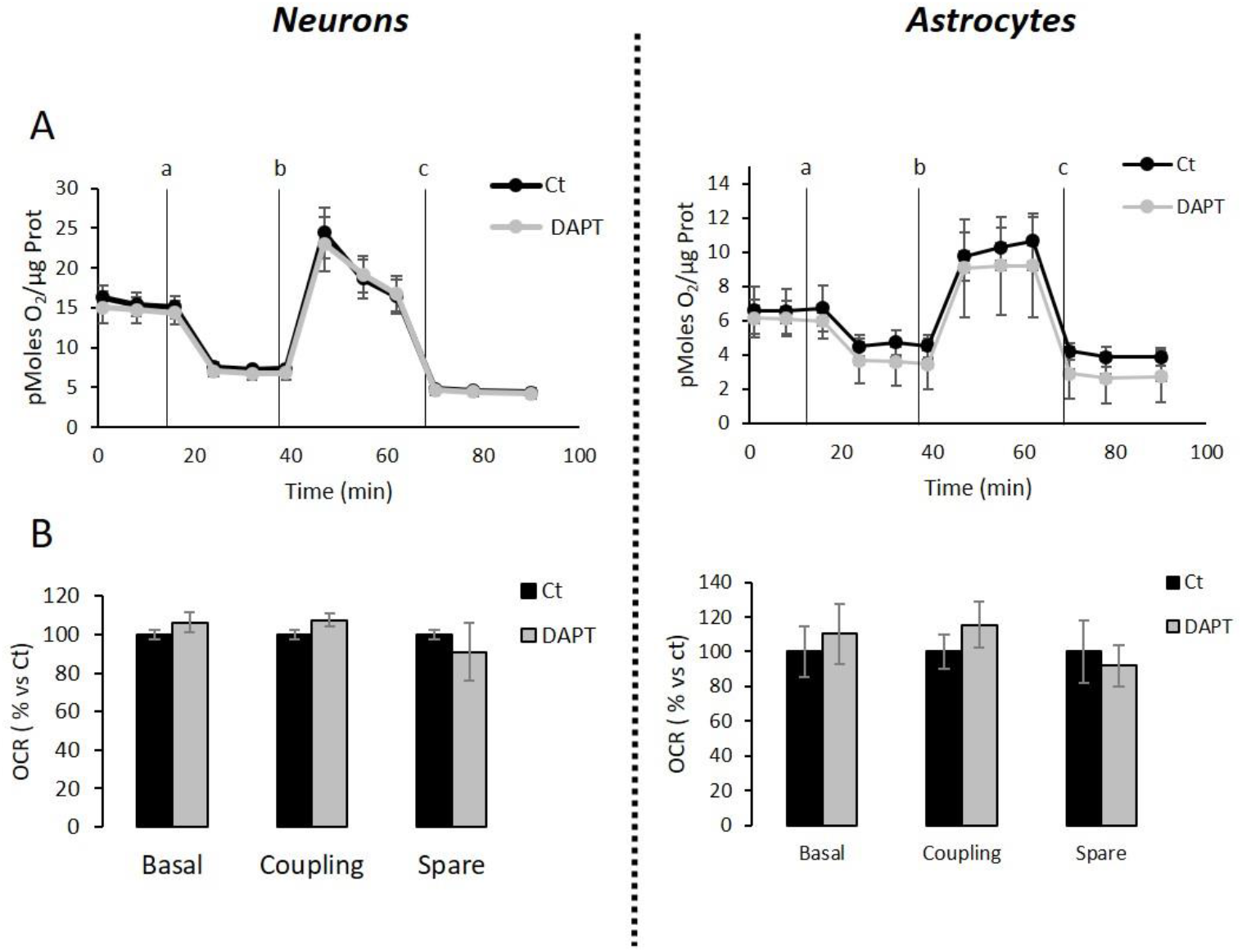
Assessment of the OXPHOS capacity in primary nerve cultures treated with DAPT. OCR was evaluated by using the Seahorse XF96 bioenergetic analyzer. Experiments were performed in primary neurons (DIV11) or primary astrocytes (DIV7) wild-type non-treated (Ct) or treated with DAPT. **(A)** General profile of the OCR with vertical lines indicate the time point at which the different compounds have been added: a. Oligomycin (CV inhibitor) b. FCCP (ΔΨ uncoupler) c. Rotenone (CI inhibitor) and antimycin A (CIII inhibitor). Values (means ± sem) are given in pmol O_2_/min/μg protein (N=3). **(B)** The basal respiration, the coupling ratio and the spare respiratory capacity were calculated according to the Cell Mito Stress Test kit’s recommended protocol. Values (means ± sem) are given as percentage of signal measured in the respective control cells (Ct or Mock). Student’s t test for neurons results (N=3) and Mann-Whitney test for astrocytes results (N = 2).

Finally, we measured the NAD^+^/NADH ratio (Figure 6A) and glycolytic flux (Figure 6B) which are parameters related to bioenergetics. NADH is an electron donor used by the first complex of the ETC. Glycolysis can either produce intermediates for OXPHOS or produce ATP and lactate depending on the oxidative status. These parameters were both decreased in PS2KO MEF cell lines. We did not observe any changes in these parameters in primary neurons or astrocytes when PSs are not expressed. In agreement, ATP levels were stable in all cell types (Figure 6C). All these data strongly support that the mitochondrial activity and related bioenergetics are not dependent on PSs in neurons and astrocytes, on the contrary to what was observed in MEF (Contino et al., 2017).

**Figure 6:**
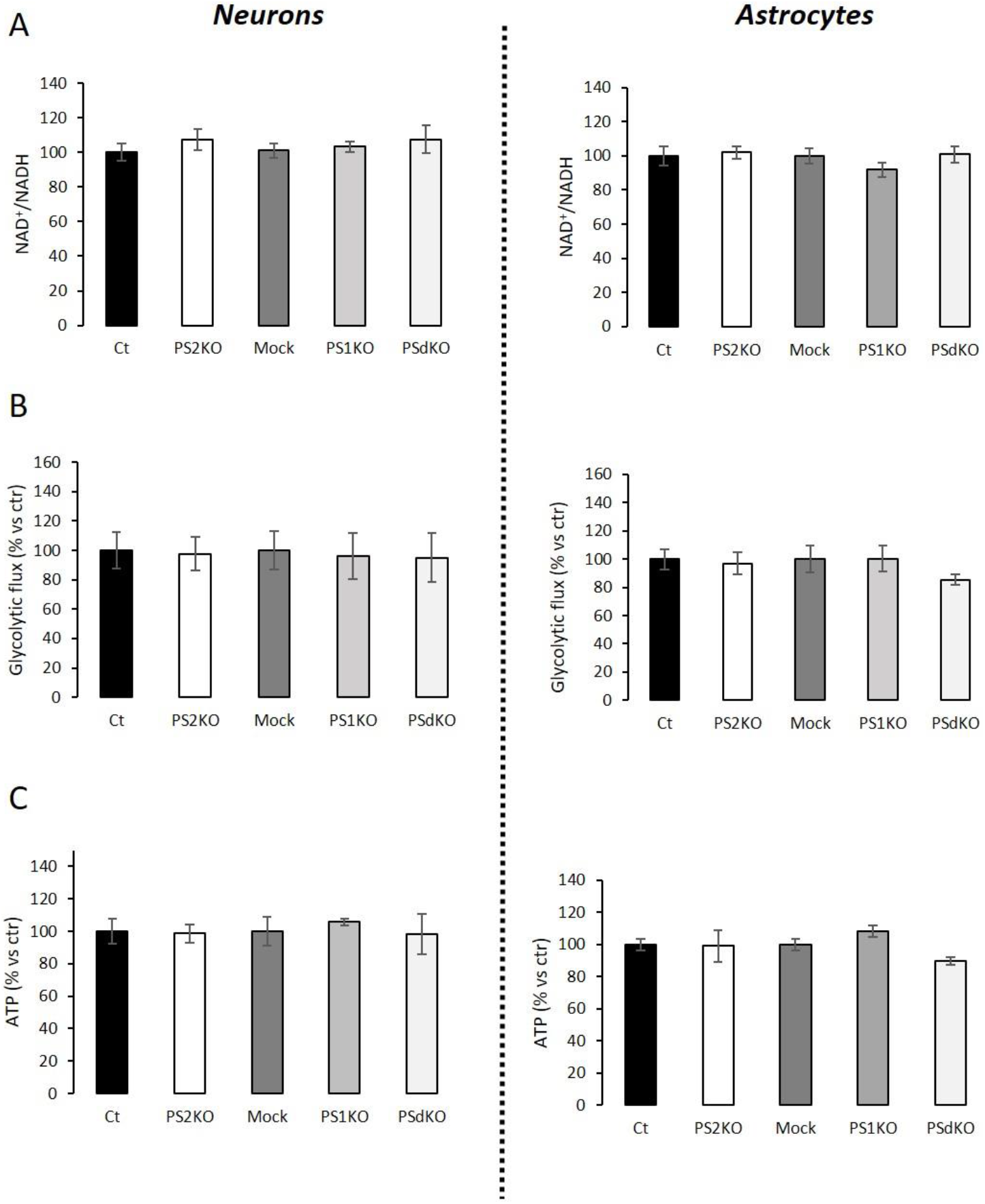
Evaluation of ATP levels, NAD^+^/NADH ratio and glycolytic flux. Experiments were performed respectively at DIV11 and DIV24 in neuronal (left panel) and astrocytes (right panel) cultures. Cell conditions were wild-type non-infected (Ct) vs PS2KO; Mock (infection control) vs PS1KO and PSdKO. (ctr= Ct or Mock) **(A)** NAD^+^/NADH ratio was quantified by a bioluminescent kit (min N=3). **(B)** Glycolysis rate was determined by the detritation rate of [3-^3^H] glucose after a 30 min incubation. Data were normalized to protein content (min N=3). **(C)** ATP level was quantified by a bioluminescent kit (min N=3). ANOVA and Tukey’s multiple comparison test.

### 3.4. Metabolic characterization of PS2 deficient primary MEF

Since no metabolic defect was observed in primary neuronal and astrocyte cells, we hypothesized that the phenotype previously observed in MEF cells could be exclusively peripheral. Indeed, the general OCR profile and related parameters were defective in PS2KO MEF cell lines and restored after stable re-expression of human PS2 (suppl. data Figure 1A). To further investigate this idea, we decided to generate primary MEF derived from E16 WT (Ct) or PS2KO mice. We measured the expression of TOM20 and subunits of OXPHOS complexes (Figure 7B) and did not observe any differences between genotypes. The activity of the ETC was also stable in the absence of PS2 as shown with the general OCR profile and the related parameters (Figure 7C). Finally, NAD+/NADH ratio and glycolytic flux (Figure 7D-E) were not modified either, indicating a metabolic stability in primary fibroblasts lacking PS2. Thus, the metabolic phenotype observed in primary fibroblasts is not consistent with the one reported in immortalized MEF.

**Figure 7:**
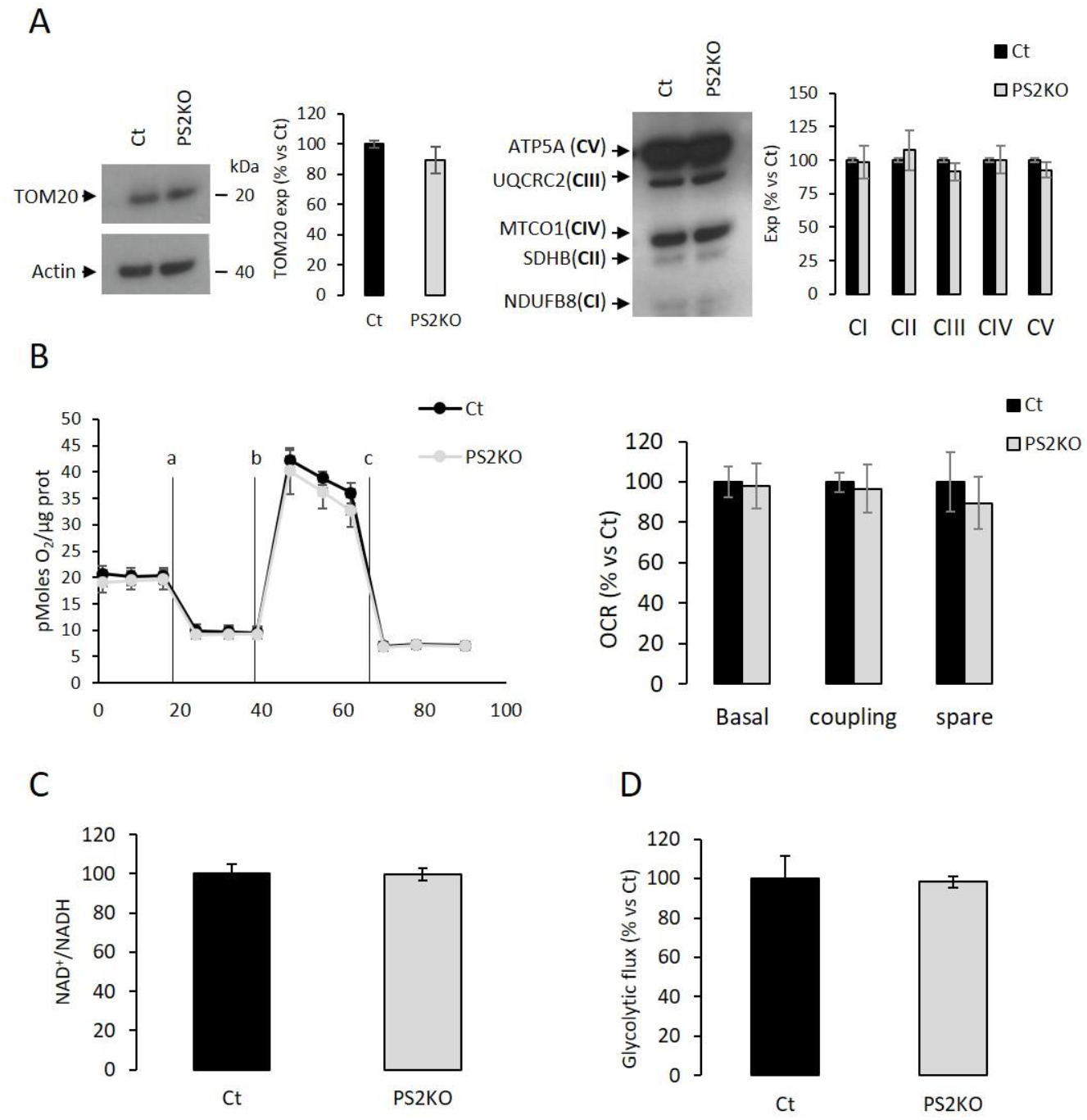
Metabolic characterization of PS2 deficient primary MEF. Experiments were carried out in primary MEF derived from E16 WT mice (Ct) or PS2KO mice (PS2KO). **(A)** Expression profile of TOM20 (left panel; N=5) and representative protein subunits from each of the five mitochondrial complexes (right panel; NDUFB8 for CI; SDHB for CII; UQCRC2 for CIII; MTCO1 for CIV; ATP5A for CV; N=5) evaluated by WB on cell lysates. Actin was used as a loading control. Quantification of the different WBs (means ± sem) are given as percentage of signal measured in the respective control cells (Ct or Mock) (min N= 3). Student’s t test. **(B)** Left panel: General OCR profile of primary MEF Ct vs PS2KO. Values (means ± sem) given in pmol O_2_/min/μg protein. Right panel: The basal respiration, the coupling ratio and the spare respiratory capacity. Values (means ± sem) are given as percentage of signal measured in the respective control cells (Ct or Mock) (N=3). **(C)** NAD^+^/NADH ratio was quantified by a bioluminescent kit (N=4). **(D)** Glycolytic rate was determined by the detritation rate of [3-3H] glucose after a 30 min incubation. Data were normalized to protein content (N=3). Student’s t test.

## 4. Discussion

PSs play a major role in cell physiology as catalytic subunits of the γ-secretase complex. γ-secretase is a multiprotein membrane complex, involved in regulated intramembrane proteolysis (RIP). Up to 90 membrane proteins have been identified as substrates of this complex (Haapasalo & Kovacs, 2011). Major substrates are Notch and the Amyloid Precursor Protein (APP), which play a critical role during development and in the amyloid pathology found in AD (Hardy, 2006). Still, in addition to membrane protein proteolysis, PSs have also been implicated in other cellular functions, including the regulation of calcium homeostasis, cell-cell adhesion, membrane trafficking, and a scaffolding role in Wnt signaling (Otto, Sharma, & Williams, 2016). The most reported non-catalytic functions of PSs are calcium homeostasis (Cheung et al., 2008; Nelson et al., 2007; Wu et al., 2013) and the building of the ER/mitochondria functional interfaces called mitochondria-associated membranes (MAMs) (Area-Gomez et al., 2009; Brunello et al., 2009; Filadi et al., 2016).

We previously observed in MEF cell lines that the absence of PS2 led to a decrease in OXPHOS activity and expression, along with an increased anaerobic glycolysis that sustains the ATP production. These effects were cell-autonomous since bioenergetic defects were rescued by the stable expression of human PS2 in PS2KO cells (Contino et al., 2017). These were strong incentives supporting the role of Presenilins, and more precisely PS2 in mitochondrial function. Several studies were in line with these observations. PS2 but not PS1 deficiency was previously reported to alter mitochondrial respiration indicating a PS2-specific role in mitochondrial function (Behbahani et al., 2006). PS2 was found to modulate calcium shuttling between the ER and mitochondria, a critical process for OXPHOS stimulation and therefore mitochondrial activity (Brunello et al., 2009). In agreement, a study reported that PSs are enriched at the junction between ER and mitochondria, in domains called MAMs (Area-Gomez et al., 2009). Most of these results were obtained in immortalized MEF or in neuronal cell lines.

We went further by investigating the role of PSs in mitochondrial function using mouse primary neurons and astrocytes. Contrary to what was reported before and what we previously observed, we found that the absence of PS1 or PS2 and even of both (PSdKO) does not affect mitochondrial ETC and related bioenergetic parameters in primary nerve cells. Importantly, we also did not observe any change in primary PS2-deficient fibroblasts, in opposition to our previous observations in immortalized fibroblasts (MEF cell lines). This raises two very important points about the use of MEF to investigate PSs functions. First, the PS2KO MEF cell lines, which have been widely used so far can genetically derive and acquire clonal properties, with the risk of artifacts or misinterpretation of the data (Kaur & Dufour, 2012). Indeed, immortalized cell lines can acquire mutations with subcultures and these mutations can interfere with the cellular phenotype. However, the rescue experiments that we performed indicated that the mitochondrial defects observed in PS2KO MEF cell lines are truly PS2-dependent (Contino et al., 2017). Second, the immortalization process specifically revealed a PS2-dependent phenotype. The fact that a PS2-dependant phenotype is observed only in immortalized cells might be supported by some pieces of evidence in the literature. PSs have been related to cancer through different pathways and the inhibition of γ-secretase activity could be a potential cancer treatment targeting Notch and Wnt signaling. Indeed, PSs clearly reulate to Notch and Wnt pathways, which are both involved in cancer development (Andersson & Lendahl, 2014; Li et al., 2016; Xia et al., 2001). An increase in lung tumor formation through peroxiredoxin 6 (PRDX6) activation was also reported in PS2KO mice. The proposed underlying mechanism supports that the PS2-dependent γ-secretase cleavage of PRDX6 inhibits its critical activity in cell growth (Park et al., 2017; Yun et al., 2014). Therefore, PS2KO mice are more sensitive to develop tumors. Another study showed that PSs might be involved in epidermal growth and transformation through regulating the epidermal growth factor (EGFR) signaling (Rocher-Ros et al., 2010). It is also important to take into account that the immortalization process used for the generation of MEF cell lines was based on SV40 Antigen T expression (De Strooper et al., 1999), which shows similarities with tumor development due to the large T antigen forming complexes with pRB-1 and p53 (Hubbard & Ozer, 1999; Pipas, 2009). Interestingly, PSs have been associated to regulation of proteins such as for Akt, HIFα or β-catenin (De Gasperi, Sosa, Dracheva, & Elder, 2010; Kang et al., 2005; Xia et al., 2001). Those proteins contribute to the Warburg effect which states that most cancer cells produce lactic acid from glucose even under non-hypoxic conditions and despite functional mitochondria (Koppenol, Bounds, & Dang, 2011). This Warburg effect might have been revealed in absence of PS2 in immortalized cells. Generation of other PS2KO cell lines would therefore be useful to understand whether the mitochondrial alterations reported in PS2KO cells appear upon immortalization. The large amount of data reported so far in these cell types (Behbahani et al., 2006; Brunello et al., 2009; Filadi et al., 2016; Pera et al., 2017; Tu et al., 2006) could be relevant to determine a role of PSs and γ-secretase activity in cancer models, but this is difficult to translate to a neurodegenerative context. For this, the use of nerve cells rather that immortalized cell lines appears necessary.

To that end, we performed our study on primary neuronal and astrocyte cultures. Neurons are a prime target of neurodegeneration, and the main function of astrocyte is to support neuronal activity. More precisely, impairment of neuronal network and activity imply gradual memory and other cognitive deficits in AD (Long & Holtzman, 2019). Although neurons are the most studied nerve cell type, astrocytes were shown to be very important in the outcome of the pathology (Verkhratsky, Olabarria, Noristani, Yeh, & Rodriguez, 2010). Recent evidence suggest that early stages of neurodegenerative processes are associated with atrophy of astroglia, which causes disruptions in synaptic connectivity and excitotoxicity. At the later stages, astrocytes become activated and contribute to neuroinflammation. Also, these two nerve cell types are known to be metabolically different with neurons relying primarily on mitochondrial oxidative phosphorylation and astrocytes on glycolysis. However, we did not observe any metabolic defects in our PS-deficient primary cultures. Indeed, the OXPHOS system (activity and expression) as well as the ΔΨ, glycolysis and NAD^+^/NADH ratio were comparable between control, PS2KO, PS1KO and PSdKO cells. Since astrocytes are more glycolytic compared to neurons (Castelli et al., 2019; Kasischke, Vishwasrao, Fisher, Zipfel, & Webb, 2004), we hypothesized that astrocytes would be close, in terms of metabolic phenotype, to the MEF cell line we previously analyzed (Contino et al., 2017). However, all the metabolic parameters evaluated were similar between genotypes and these data lead us to conclude that the lack of PSs does not affect neither the OXPHOS, nor related metabolic aspects, such as glycolysis and NAD^+^/NADH ratio, in primary nerve cells. Interestingly, similar discrepancies have been already observed in a metabolic study comparing primary neurons, astrocytes and fibroblasts cultures deficient for the mitochondrial complex I subunit Ndufs4, a model for the mitochondrial Leigh Syndrome disease (Bird et al., 2014). The ΔΨ and ATP synthesis were impaired in the NDUFS4 KO primary MEF, with an increase of ROS generation and an altered sensitivity to cell death. In contrast, NDUFS4 KO primary neurons and astrocytes displayed only impaired ATP generation. This observation is of high interest since Leigh Syndrome is a neurological pathology and one could expect to observe the defects in nerve cultures and not in fibroblasts. In this study (Bird et al., 2014), the primary MEF were not immortalized but revealed how differences can be observed from one cell type to the other, in particular between central and peripheral cells.

Still, the fact that the absence of PSs does not affect metabolism is rather intriguing. Considering the expression profiles of the *PSEN1* and *PSEN2* genes throughout the body, one could expect to have distinct phenotypes related either to PS1 or PS2 deficiency. Indeed, PS1 was suggested to be more important in CNS and PS2 in peripheral organs (Lee et al., 1996). However, even PS1KO nerve cells were not metabolically defective. The PSdKO cells also exhibited no defects, which was surprisingly unexpected considering all the attributed functions to PSs (Wolfe, 2019; Zhang et al., 2013), and that the complex of which they are part of cleaves more than 90 substrates (Haapasalo & Kovacs, 2011; Wolfe, 2019). Especially, cortical brain sections from conditional PSdKO mice show several pathological features such as neurodegeneration, astrogliosis and swollen mitochondria (Saura et al., 2004; Wines-Samuelson et al., 2010). Importantly, the experimental set-up and environment should be taken into consideration to give a correct interpretation related to bioenergetics. All our experiments were performed on primary cells that were cultured in low glucose (5 mM) medium, to reflect a more physiological condition, that were mature and differentiated (DIV11 for neurons and DIV24 for astrocytes) and that were at basal and resting state. Challenging the cells with depolarizing conditions or hypoxia could perhaps unravel a PS-dependent phenotype. Moreover, we performed relative pure cultures of one specific cell type that does not reflect cell-to-cell communication and exchanges happening in the CNS context. This challenging multicellular context might explain why in PSdKO mice, mitochondria seem affected. Indeed, in that case, animal models constitute much more complex systems than *in vitro* cultures bearing different cell types, cell interactions and processes that were found to be disrupted in absence of PSs (Saura et al., 2004; Wines-Samuelson et al., 2010).

In conclusion, our study provides new evidence for the lack of mitochondrial alterations in PS1KO, PS2KO and even PSdKO neurons and astrocytes, as well as in PS2KO primary MEF. This is contradictory to previous observations in neuronal cell lines and immortalized fibroblasts. Thus, immortalized cells might provide relevant results for the role of PSs in mitochondrial activity and bioenergetics related to cancer processes, in which PSs involvement have already been reported. However, the contribution of PSs to alterations in mitochondrial activity related to neurodegenerative processes, such as AD, needs to be critically readdressed and further explored in relevant models.

## Abbreviations

ΔΨ: mitochondrial membrane potential
2R2: PS2 knockout rescued PS2
AD: Alzheimer’s disease
CI-CV: complexes I-V
Ct: control
DTAB: dodecyltrimethylammonium bromide
ETC: electron transport chain
ER: endoplasmic reticulum
FAD: familial AD
FCCP: carbonyl cyanide-4-(trifluoromethoxy) phenylhydrazone
GFAP: glial-fibrillary acidic protein
MAM: mitochondria-associated membranes
MAP-2: microtubule-associated protein 2
MEF: mouse embryonic fibroblasts
OCR: oxygen consumption rate
OXPHOS: oxidative phosphorylation
PS: presenilin
ROS: reactive oxygen species
TMRM: tetramethylrhodamine methyl ester
WB: western blotting.

---

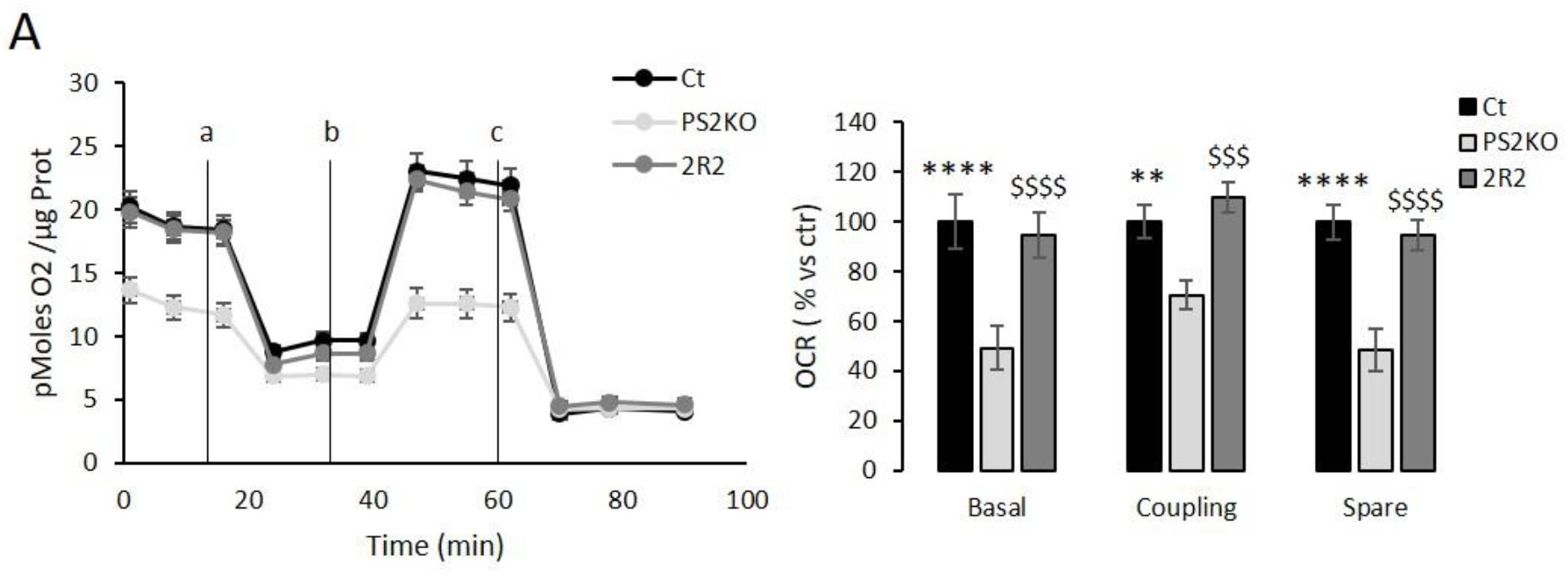
Supplemental data: Oxygen consumption profile in immortalized PS2 deficient MEF cell line. **(A)** General OCR profile of immortalized MEF, wild type (Ct), PS2KO and PS2KO rescued PS2 (2R2). Vertical lines indicate the time point at which the different compounds have been added: a. Oligomycin (CV inhibitor) b. FCCP (ΔΨ uncoupler) c. Rotenone (CI inhibitor) and antimycin A (CIII inhibitor). Values (means ± sem) are given in pmol O_2_/min/μg protein. Right panel: The basal respiration, the coupling ratio and the spare respiratory capacity were calculated according to the Cell Mito Stress Test kit’s recommended protocol. Values (means ± sem) are given **as percentage of signal measured in the Ct. * vs Ct and $ vs 2R2. ****p < 0.0001; **p= 0.0022.** ANOVA and Tukey’s multiple comparison test (N=1; n=6).

## Author credit statement

SC performed research, analyzed data and wrote the paper. C.V. and N.S. performed research and analyzed data. D.V. performed research V.P., S.S. and L.B. analyzed data. P.K.C. designed research and wrote the paper.

## Acknowledgements

L.B. is research Director of FRS-FNRS. S.C. is supported by a fellowship “hope in head” from the Rotary. C.V is supported by a fellowship from the Belgian Fonds pour la formation à la Recherche dans l’Industrie et l’Agriculture (FRIA-FNRS). We thank Bernadette Tasiaux for technical support.

## References

Adav, S. S., Park, J. E., & Sze, S. K. (2019). Quantitative profiling brain proteomes revealed mitochondrial dysfunction in Alzheimer’s disease. Mol Brain, 12 (1), 8. doi:10.1186/s13041-019-0430-y

Andersson, E. R., & Lendahl, U. (2014). Therapeutic modulation of Notch signalling--are we there yet? Nat Rev Drug Discov, 13 (5), 357–378. doi:10.1038/nrd4252

Area-Gomez, E., de Groof, A. J., Boldogh, I., Bird, T. D., Gibson, G. E., Koehler, C. M., … Schon, E. A. (2009). Presenilins are enriched in endoplasmic reticulum membranes associated with mitochondria. Am J Pathol, 175 (5), 1810–1816. doi:10.2353/ajpath.2009.090219

Behbahani, H., Shabalina, I. G., Wiehager, B., Concha, H., Hultenby, K., Petrovic, N., … Ankarcrona, M. (2006). Differential role of Presenilin-1 and −2 on mitochondrial membrane potential and oxygen consumption in mouse embryonic fibroblasts. J Neurosci Res, 84 (4), 891–902. doi:10.1002/jnr.20990

Bird, M. J., Wijeyeratne, X. W., Komen, J. C., Laskowski, A., Ryan, M. T., Thorburn, D. R., & Frazier, A. E. (2014). Neuronal and astrocyte dysfunction diverges from embryonic fibroblasts in the Ndufs4fky/fky mouse. Biosci Rep, 34 (6), e00151. doi:10.1042/BSR20140151

Brunello, L., Zampese, E., Florean, C., Pozzan, T., Pizzo, P., & Fasolato, C. (2009). Presenilin-2 dampens intracellular Ca2^+^ stores by increasing Ca2^+^ leakage and reducing Ca2^+^ uptake. J Cell Mol Med, 13 (9B), 3358–3369. doi:10.1111/j.1582-4934.2009.00755.x

Castelli, V., Benedetti, E., Antonosante, A., Catanesi, M., Pitari, G., Ippoliti, R., … d’Angelo, M. (2019). Neuronal Cells Rearrangement During Aging and Neurodegenerative Disease: Metabolism, Oxidative Stress and Organelles Dynamic. Front Mol Neurosci, 12, 132. doi:10.3389/fnmol.2019.00132

Ceyzeriat, K., Zilli, T., Millet, P., Frisoni, G. B., Garibotto, V., & Tournier, B. B. (2020). Learning from the Past: A Review of Clinical Trials Targeting Amyloid, Tau and Neuroinflammation in Alzheimer’s Disease. Curr Alzheimer Res, 17 (2), 112–125. doi:10.2174/1567205017666200304085513

Charlesworth, P., Cotterill, E., Morton, A., Grant, S. G., & Eglen, S. J. (2015). Quantitative differences in developmental profiles of spontaneous activity in cortical and hippocampal cultures. Neural Dev, 10, 1. doi:10.1186/s13064-014-0028-0

Cheung, K. H., Shineman, D., Muller, M., Cardenas, C., Mei, L., Yang, J., … Foskett, J. K. (2008). Mechanism of Ca2^+^ disruption in Alzheimer’s disease by presenilin regulation of InsP3 receptor channel gating. Neuron, 58 (6), 871–883. doi:10.1016/j.neuron.2008.04.015

Contino, S., Porporato, P. E., Bird, M., Marinangeli, C., Opsomer, R., Sonveaux, P., … Kienlen-Campard, P. (2017). Presenilin 2-Dependent Maintenance of Mitochondrial Oxidative Capacity and Morphology. Front Physiol, 8, 796. doi:10.3389/fphys.2017.00796

Correia, S. C., Perry, G., & Moreira, P. I. (2016). Mitochondrial traffic jams in Alzheimer’s disease - pinpointing the roadblocks. Biochim Biophys Acta, 1862 (10), 1909–1917. doi:10.1016/j.bbadis.2016.07.010

De Gasperi, R., Sosa, M. A., Dracheva, S., & Elder, G. A. (2010). Presenilin-1 regulates induction of hypoxia inducible factor-1alpha: altered activation by a mutation associated with familial Alzheimer’s disease. Mol Neurodegener, 5, 38. doi:10.1186/1750-1326-5-38

De Strooper, B., Annaert, W., Cupers, P., Saftig, P., Craessaerts, K., Mumm, J. S., … Kopan, R. (1999). A presenilin-1-dependent gamma-secretase-like protease mediates release of Notch intracellular domain. Nature, 398 (6727), 518–522. doi:10.1038/19083

Dixit, S., Fessel, J. P., & Harrison, F. E. (2017). Mitochondrial dysfunction in the APP/PSEN1 mouse model of Alzheimer’s disease and a novel protective role for ascorbate. Free Radic Biol Med, 112, 515–523. doi:10.1016/j.freeradbiomed.2017.08.021

Donoviel, D. B., Hadjantonakis, A. K., Ikeda, M., Zheng, H., Hyslop, P. S., & Bernstein, A. (1999). Mice lacking both presenilin genes exhibit early embryonic patterning defects. Genes Dev, 13 (21), 2801–2810. doi:10.1101/gad.13.21.2801

El-Brolosy, M. A., & Stainier, D. Y. R. (2017). Genetic compensation: A phenomenon in search of mechanisms. PLoS Genet, 13 (7), e1006780. doi:10.1371/journal.pgen.1006780

Filadi, R., Greotti, E., Turacchio, G., Luini, A., Pozzan, T., & Pizzo, P. (2016). Presenilin 2 Modulates Endoplasmic Reticulum-Mitochondria Coupling by Tuning the Antagonistic Effect of Mitofusin 2. Cell Rep, 15 (10), 2226–2238. doi:10.1016/j.celrep.2016.05.013

Filadi, R., Theurey, P., & Pizzo, P. (2017). The endoplasmic reticulum-mitochondria coupling in health and disease: Molecules, functions and significance. Cell Calcium, 62, 1–15. doi:10.1016/j.ceca.2017.01.003

Garcia-Escudero, V., Martin-Maestro, P., Perry, G., & Avila, J. (2013). Deconstructing mitochondrial dysfunction in Alzheimer disease. Oxid Med Cell Longev, 2013, 162152. doi:10.1155/2013/162152

Goate, A., Chartier-Harlin, M. C., Mullan, M., Brown, J., Crawford, F., Fidani, L., … et al. (1991). Segregation of a missense mutation in the amyloid precursor protein gene with familial Alzheimer’s disease. Nature, 349 (6311), 704–706. doi:10.1038/349704a0

Green, K. N., Demuro, A., Akbari, Y., Hitt, B. D., Smith, I. F., Parker, I., & LaFerla, F. M. (2008). SERCA pump activity is physiologically regulated by presenilin and regulates amyloid beta production. J Cell Biol, 181 (7), 1107–1116. doi:10.1083/jcb.200706171

Haapasalo, A., & Kovacs, D. M. (2011). The many substrates of presenilin/gamma-secretase. J Alzheimers Dis, 25 (1), 3–28. doi:10.3233/JAD-2011-101065

Hardy, J. (2006). Alzheimer’s disease: the amyloid cascade hypothesis: an update and reappraisal. J Alzheimers Dis, 9(3 Suppl), 151–153. doi:10.3233/jad-2006-9s317

Herholz, K. (2012). Use of FDG PET as an imaging biomarker in clinical trials of Alzheimer’s disease. Biomark Med, 6 (4), 431–439. doi:10.2217/bmm.12.51

Herreman, A., Hartmann, D., Annaert, W., Saftig, P., Craessaerts, K., Serneels, L., … De Strooper, B. (1999). Presenilin 2 deficiency causes a mild pulmonary phenotype and no changes in amyloid precursor protein processing but enhances the embryonic lethal phenotype of presenilin 1 deficiency. Proc Natl Acad Sci U S A, 96 (21), 11872–11877. doi:10.1073/pnas.96.21.11872

Hubbard, K., & Ozer, H. L. (1999). Mechanism of immortalization. Age (Omaha), 22 (2), 65–69. doi:10.1007/s11357-999-0008-1

Kang, D. E., Yoon, I. S., Repetto, E., Busse, T., Yermian, N., Ie, L., & Koo, E. H. (2005). Presenilins mediate phosphatidylinositol 3-kinase/AKT and ERK activation via select signaling receptors. Selectivity of PS2 in platelet-derived growth factor signaling. J Biol Chem, 280 (36), 31537–31547. doi:10.1074/jbc.M500833200

Kasischke, K. A., Vishwasrao, H. D., Fisher, P. J., Zipfel, W. R., & Webb, W. W. (2004). Neural activity triggers neuronal oxidative metabolism followed by astrocytic glycolysis. Science, 305 (5680), 99–103. doi:10.1126/science.1096485

Kaur, G., & Dufour, J. M. (2012). Cell lines: Valuable tools or useless artifacts. Spermatogenesis, 2 (1), 1–5. doi:10.4161/spmg.19885

Koppenol, W. H., Bounds, P. L., & Dang, C. V. (2011). Otto Warburg’s contributions to current concepts of cancer metabolism. Nat Rev Cancer, 11 (5), 325–337. doi:10.1038/nrc3038

Lee, M. K., Slunt, H. H., Martin, L. J., Thinakaran, G., Kim, G., Gandy, S. E., … Sisodia, S. S. (1996). Expression of presenilin 1 and 2 (PS1 and PS2) in human and murine tissues. J Neurosci, 16 (23), 7513–7525.

Li, P., Lin, X., Zhang, J. R., Li, Y., Lu, J., Huang, F. C., … Huang, C. M. (2016). The expression of presenilin 1 enhances carcinogenesis and metastasis in gastric cancer. Oncotarget, 7 (9), 10650–10662. doi:10.18632/oncotarget.7298

Long, J. M., & Holtzman, D. M. (2019). Alzheimer Disease: An Update on Pathobiology and Treatment Strategies. Cell, 179 (2), 312–339. doi:10.1016/j.cell.2019.09.001

Marsin, A. S., Bouzin, C., Bertrand, L., & Hue, L. (2002). The stimulation of glycolysis by hypoxia in activated monocytes is mediated by AMP-activated protein kinase and inducible 6-phosphofructo-2-kinase. J Biol Chem, 277 (34), 30778–30783. doi:10.1074/jbc.M205213200

Martin-Maestro, P., Gargini, R., Garcia, E., Perry, G., Avila, J., & Garcia-Escudero, V. (2017). Slower Dynamics and Aged Mitochondria in Sporadic Alzheimer’s Disease. Oxid Med Cell Longev, 2017, 9302761. doi:10.1155/2017/9302761

Mosconi, L., Berti, V., Glodzik, L., Pupi, A., De Santi, S., & de Leon, M. J. (2010). Pre-clinical detection of Alzheimer’s disease using FDG-PET, with or without amyloid imaging. J Alzheimers Dis, 20 (3), 843–854. doi:10.3233/JAD-2010-091504

Nelson, O., Tu, H., Lei, T., Bentahir, M., de Strooper, B., & Bezprozvanny, I. (2007). Familial Alzheimer disease-linked mutations specifically disrupt Ca2^+^ leak function of presenilin 1. J Clin Invest, 117 (5), 1230–1239. doi:10.1172/JCI30447

Opsomer, R., Contino, S., Perrin, F., Gualdani, R., Tasiaux, B., Doyen, P., … Kienlen-Campard, P. (2020). Amyloid Precursor Protein (APP) Controls the Expression of the Transcriptional Activator Neuronal PAS Domain Protein 4 (NPAS4) and Synaptic GABA Release. eNeuro, 7(3). doi:10.1523/ENEURO.0322-19.2020

Otto, G. P., Sharma, D., & Williams, R. S. (2016). Non-Catalytic Roles of Presenilin Throughout Evolution. J Alzheimers Dis, 52 (4), 1177–1187. doi:10.3233/JAD-150940

Pagani, L., & Eckert, A. (2011). Amyloid-Beta interaction with mitochondria. Int J Alzheimers Dis, 2011, 925050. doi:10.4061/2011/925050

Park, M. H., Yun, H. M., Hwang, C. J., Park, S. I., Han, S. B., Hwang, D. Y., … Hong, J. T. (2017). Presenilin Mutation Suppresses Lung Tumorigenesis via Inhibition of Peroxiredoxin 6 Activity and Expression. Theranostics, 7 (15), 3624–3637. doi:10.7150/thno.21408

Pera, M., Larrea, D., Guardia-Laguarta, C., Montesinos, J., Velasco, K. R., Agrawal, R. R., … Area-Gomez, E. (2017). Increased localization of APP-C99 in mitochondria-associated ER membranes causes mitochondrial dysfunction in Alzheimer disease. EMBO J, 36 (22), 3356–3371. doi:10.15252/embj.201796797

Pipas, J. M. (2009). SV40: Cell transformation and tumorigenesis. Virology, 384 (2), 294–303. doi:10.1016/j.virol.2008.11.024

Reddy, P. H. (2013). Amyloid beta-induced glycogen synthase kinase 3beta phosphorylated VDAC1 in Alzheimer’s disease: implications for synaptic dysfunction and neuronal damage. Biochim Biophys Acta, 1832 (12), 1913–1921. doi:10.1016/j.bbadis.2013.06.012

Rocher-Ros, V., Marco, S., Mao, J. H., Gines, S., Metzger, D., Chambon, P., … Saura, C. A. (2010). Presenilin modulates EGFR signaling and cell transformation by regulating the ubiquitin ligase Fbw7. Oncogene, 29 (20), 2950–2961. doi:10.1038/onc.2010.57

Rogaev, E. I., Sherrington, R., Rogaeva, E. A., Levesque, G., Ikeda, M., Liang, Y., … et al. (1995). Familial Alzheimer’s disease in kindreds with missense mutations in a gene on chromosome 1 related to the Alzheimer’s disease type 3 gene. Nature, 376 (6543), 775–778. doi:10.1038/376775a0

Saura, C. A., Choi, S. Y., Beglopoulos, V., Malkani, S., Zhang, D., Shankaranarayana Rao, B. S., … Shen, J. (2004). Loss of presenilin function causes impairments of memory and synaptic plasticity followed by age-dependent neurodegeneration. Neuron, 42 (1), 23–36. doi:10.1016/s0896-6273(04)00182-5

Schildge, S., Bohrer, C., Beck, K., & Schachtrup, C. (2013). Isolation and culture of mouse cortical astrocytes. J Vis Exp(71). doi:10.3791/50079

Serrano-Pozo, A., Frosch, M. P., Masliah, E., & Hyman, B. T. (2011). Neuropathological alterations in Alzheimer disease. Cold Spring Harb Perspect Med, 1 (1), a006189. doi:10.1101/cshperspect.a006189

Shen, J., Bronson, R. T., Chen, D. F., Xia, W., Selkoe, D. J., & Tonegawa, S. (1997). Skeletal and CNS defects in Presenilin-1-deficient mice. Cell, 89 (4), 629–639. doi:10.1016/s0092-8674(00)80244-5

Sherrington, R., Rogaev, E. I., Liang, Y., Rogaeva, E. A., Levesque, G., Ikeda, M., … St George-Hyslop, P. H. (1995). Cloning of a gene bearing missense mutations in early-onset familial Alzheimer’s disease. Nature, 375 (6534), 754–760. doi:10.1038/375754a0

Shivamurthy, V. K., Tahari, A. K., Marcus, C., & Subramaniam, R. M. (2015). Brain FDG PET and the diagnosis of dementia. AJR Am J Roentgenol, 204 (1), W76–85. doi:10.2214/AJR.13.12363

Stanga, S., Caretto, A., Boido, M., & Vercelli, A. (2020). Mitochondrial Dysfunctions: A Red Thread across Neurodegenerative Diseases. Int J Mol Sci, 21(10). doi:10.3390/ijms21103719

Stanga, S., Vrancx, C., Tasiaux, B., Marinangeli, C., Karlstrom, H., & Kienlen-Campard, P. (2018). Specificity of presenilin-1- and presenilin-2-dependent gamma-secretases towards substrate processing. J Cell Mol Med, 22 (2), 823–833. doi:10.1111/jcmm.13364

Stanga, S., Zanou, N., Audouard, E., Tasiaux, B., Contino, S., Vandermeulen, G., … Kienlen-Campard, P. (2016). APP-dependent glial cell line-derived neurotrophic factor gene expression drives neuromuscular junction formation. FASEB J, 30 (5), 1696–1711. doi:10.1096/fj.15-278739

Swerdlow, R. H. (2018). Mitochondria and Mitochondrial Cascades in Alzheimer’s Disease. J Alzheimers Dis, 62 (3), 1403–1416. doi:10.3233/JAD-170585

Swerdlow, R. H., Burns, J. M., & Khan, S. M. (2014). The Alzheimer’s disease mitochondrial cascade hypothesis: progress and perspectives. Biochim Biophys Acta, 1842 (8), 1219–1231. doi:10.1016/j.bbadis.2013.09.010

Tu, H., Nelson, O., Bezprozvanny, A., Wang, Z., Lee, S. F., Hao, Y. H., … Bezprozvanny, I. (2006). Presenilins form ER Ca2^+^ leak channels, a function disrupted by familial Alzheimer’s disease-linked mutations. Cell, 126 (5), 981–993. doi:10.1016/j.cell.2006.06.059

Uechi, Y., Yoshioka, H., Morikawa, D., & Ohta, Y. (2006). Stability of membrane potential in heart mitochondria: single mitochondrion imaging. Biochem Biophys Res Commun, 344 (4), 1094–1101. doi:10.1016/j.bbrc.2006.03.233

Verkhratsky, A., Olabarria, M., Noristani, H. N., Yeh, C. Y., & Rodriguez, J. J. (2010). Astrocytes in Alzheimer’s disease. Neurotherapeutics, 7 (4), 399–412. doi:10.1016/j.nurt.2010.05.017

Vetrivel, K. S., Zhang, Y. W., Xu, H., & Thinakaran, G. (2006). Pathological and physiological functions of presenilins. Mol Neurodegener, 1, 4. doi:10.1186/1750-1326-1-4

Wang, X., Su, B., Lee, H. G., Li, X., Perry, G., Smith, M. A., & Zhu, X. (2009). Impaired balance of mitochondrial fission and fusion in Alzheimer’s disease. J Neurosci, 29 (28), 9090–9103. doi:10.1523/JNEUROSCI.1357-09.2009

Whitaker-Menezes, D., Martinez-Outschoorn, U. E., Flomenberg, N., Birbe, R. C., Witkiewicz, A. K., Howell, A., … Sotgia, F. (2011). Hyperactivation of oxidative mitochondrial metabolism in epithelial cancer cells in situ: visualizing the therapeutic effects of metformin in tumor tissue. Cell Cycle, 10 (23), 4047–4064. doi:10.4161/cc.10.23.18151

Wines-Samuelson, M., Schulte, E. C., Smith, M. J., Aoki, C., Liu, X., Kelleher, R. J., 3rd, & Shen, J. (2010). Characterization of age-dependent and progressive cortical neuronal degeneration in presenilin conditional mutant mice. PLoS One, 5 (4), e10195. doi:10.1371/journal.pone.0010195

Wolfe, M. S. (2019). Structure and Function of the gamma-Secretase Complex. Biochemistry, 58 (27), 2953–2966. doi:10.1021/acs.biochem.9b00401

Wu, B., Yamaguchi, H., Lai, F. A., & Shen, J. (2013). Presenilins regulate calcium homeostasis and presynaptic function via ryanodine receptors in hippocampal neurons. Proc Natl Acad Sci U S A, 110 (37), 15091–15096. doi:10.1073/pnas.1304171110

Xia, X., Qian, S., Soriano, S., Wu, Y., Fletcher, A. M., Wang, X. J., … Zheng, H. (2001). Loss of presenilin 1 is associated with enhanced beta-catenin signaling and skin tumorigenesis. Proc Natl Acad Sci U S A, 98 (19), 10863–10868. doi:10.1073/pnas.191284198

Yao, J., Irwin, R. W., Zhao, L., Nilsen, J., Hamilton, R. T., & Brinton, R. D. (2009). Mitochondrial bioenergetic deficit precedes Alzheimer’s pathology in female mouse model of Alzheimer’s disease. Proc Natl Acad Sci U S A, 106 (34), 14670–14675. doi:10.1073/pnas.0903563106

Yu, H., Kessler, J., & Shen, J. (2000). Heterogeneous populations of ES cells in the generation of a floxed Presenilin-1 allele. Genesis, 26 (1), 5–8.

Yun, H. M., Park, M. H., Kim, D. H., Ahn, Y. J., Park, K. R., Kim, T. M., … Hong, J. T. (2014). Loss of presenilin 2 is associated with increased iPLA2 activity and lung tumor development. Oncogene, 33 (44), 5193–5200. doi:10.1038/onc.2014.128

Zampese, E., Fasolato, C., Kipanyula, M. J., Bortolozzi, M., Pozzan, T., & Pizzo, P. (2011). Presenilin 2 modulates endoplasmic reticulum (ER)-mitochondria interactions and Ca2^+^ cross-talk. Proc Natl Acad Sci U S A, 108 (7), 2777–2782. doi:10.1073/pnas.1100735108

Zhang, S., Zhang, M., Cai, F., & Song, W. (2013). Biological function of Presenilin and its role in AD pathogenesis. Transl Neurodegener, 2 (1), 15. doi:10.1186/2047-9158-2-15

